# A rapidly evolved shift in life history timing during ecological speciation is driven by the transition between developmental phases

**DOI:** 10.1101/2020.01.29.925057

**Authors:** Thomas H. Q. Powell, Andrew Nguyen, Qinwen Xia, Jeffrey L. Feder, Gregory J. Ragland, Daniel A. Hahn

## Abstract

For insect species in temperate environments, seasonal timing is often governed by the regulation of diapause, a complex developmental program that allows insects to weather unfavorable conditions and synchronize their lifecycles with available resources. Diapause consists of a series of phases that govern initiation, maintenance, and termination of this developmental pathway. The evolution of insect seasonal timing depends in part on how these phases of diapause development and post-diapause development interact to affect variation in phenology. Here, we dissect the physiological basis of a recently evolved phenological shift in *Rhagoletis pomonella* (Diptera: Tephritidae), a model system for ecological divergence. A recently derived population of *R. pomonella* shifted from specializing on native hawthorn fruit to earlier fruiting introduced apples, resulting in a 3-4 week shift in adult emergence timing. We tracked metabolic rates of individual flies across post-winter development to test which phases of development may act either independently or in combination to contribute to this recently evolved divergence in timing. Apple and hawthorn flies differed in a number of facets of their post-winter developmental trajectories. However, divergent adaptation in adult emergence phenology in these flies was due almost entirely to the end of the pupal diapause maintenance phase, with post-diapause development having a very small effect. The relatively simple underpinnings of variation in adult emergence phenology suggest that further adaptation to seasonal change in these flies for this trait might be largely due to the timing of diapause termination unhindered by strong covariance among different components of post-diapause development.

**Data accessibility:** All data (in the form of tables of all metabolic rate measurements for all individual flies in the study) will be available on DRYAD when the manuscript is published.

## INTRODUCTION

Phenological adaptation can play an important role in both the origin and maintenance of biodiversity in seasonal environments. Adaptation to temporally predictable environmental variation may be a potent driver of reproductive isolation during ecological speciation (Taylor and Friesen 2017). Moreover, altered seasonality is a critical consequence of global climate change for many ecological communities (Miller-Rushing et al. 2010). Thus, understanding how evolution shapes the regulation of seasonal timing has important implications for the tempo of ecological diversification, as well as the potential of biotic communities to persist in the face of global change.

Divergent adaptation to novel habitats or niches can generate reproductive isolation between populations through the process of ecological speciation (Schluter 2001; 2009; Rundle & Nosil 2005). Temporal isolation, or allochrony, may be a common component of ecological barriers to gene flow for two reasons. First, in temperate habitats, temporally distinct resource opportunities may pose distinct trade-offs, favoring phenological specialization. Second, for organisms with seasonal reproductive biology, phenological divergence may serve as a ‘magic trait’ for speciation by pleiotropically coupling ecological divergence to assortative mating (Servedio et al. 2011). Indeed, adaptation to differences in seasonal timing has been shown to contribute to reproductive isolation in a wide variety of organisms including plants (Salvolainen et al. 2006; Lowry et al. 2008), insects (Tauber et al. 1977; Wood & Guttman 1982; Horner et al. 1999; Eubanks et al. 2003; Abbot & Withgott 2004; Ording et al. 2010; Wadsworth et al. 2013; Hood et al. 2015), and vertebrates (Friesen et al. 2007).

Shifts in phenology have also received increasing attention as anthropogenic change causes alterations in the phenology of plant and animal populations (Parmesan 2006; Van Dyck et al. 2015; Thackeray et al. 2016). Many recent studies provide evidence for fitness consequences of disrupting seasonal phenologies, from high bird nestling mortality due to mismatch with seasonal food supplies (Sanz et al. 2003) to declines in recruitment in plant populations resulting from lack of synchrony between flowering and the activity of pollinators (Rafferty & Ives 2011; Gilman et al. 2012; Kudo & Ida 2013). Cases of recent shifts in phenology during incipient speciation also provide useful windows into how the regulation of seasonal timing evolves more generally, offering insight into the factors that may promote or constrain evolutionary responses to anthropogenic change (Scriber et al. 2014, Taylor and Friesen 2017).

Phenological adaptation is crucial for species with specialist life history strategies. For example, univoltine phytophagous insects typically have one chance to breed per year, with most of these species specializing on a single host plant (Strong et al. 1984). Therefore, suboptimal timing can have severe fitness costs (Miller-Rushing et al. 2010). Many univoltine insects use seasonal dormancy, or diapause, to synchronize their active times for growth and reproduction with seasonal resources. Thus, the decision of when to enter dormancy relative to the timing of unfavorable seasonal conditions, such as the onset of winter, the dry season, or lack of host-plant availability, as well as when to exit dormancy to synchronize with favorable conditions are critical aspects of life history timing (Tauber and Tauber 1981). Phenological mismatches between insect life histories and resource availability therefore lead to low reproductive success and evolutionary dead ends (Van Dyck et al. 2015).

Insect diapause is not just a simple cessation of development resulting directly from physiologically stressful conditions (e.g., cold-induced quiescence) (Denlinger 2002; Hahn and Denlinger 2011). Rather, diapause is a dynamic, environmentally sensitive developmental program, consisting of multiple developmental phases: preparation, maintenance, and the end of diapause followed by the initiation of post-diapause development (Kostal et al. 2006). Each of these developmental phases is characterized by distinct molecular (Kostal et al. 2017; Ragland et al. 2017), morphological (Bryon et al. 2017), and physiological (Ragland et al. 2009) hallmarks. Selection to produce alternative or divergent phenological strategies may act on the relative duration of or developmental rates during these phases (Andrewartha 1952, Kostal 2006, Kostal et al. 2017). Whether or not variation in each phase of the diapause and post-diapause program represents an independent functional module underlain by distinct genetic architecture or are highly interrelated and covary has important consequences for the evolvability of seasonal timing (Wagner et al. 2007; Wagner & Zhang 2011). If independent, then it is possible that each module can be differentially shaped by natural selection to varying environmental conditions at specific points during development. However, if the timing of different phases of diapause development are highly interrelated, then it may be that selection acts on the phase having greatest significance for matching an insect’s life cycle with resource availability, with possible pleiotropic consequences on other phases. While the concept of distinct phases within the diapause developmental trajectory has long been recognized (Andrewartha 1952; Tauber et al. 1986; Kostal 2006), how different phases of diapause development interact and how selection acts on variation within these phases to affect life history timing is not well understood and has received very little attention despite its potential importance for seasonal adaptation.

To address the question of the flexibility of the diapause program to compartmentally adapt to environmental challenges requires first characterizing the different phases of diapause development in an insect and then determining their interactions (correlational structure) to assess whether they are assembled into independent modules or not. Under the simplest scenario, seasonal divergence in emergence timing is due principally to the degree of metabolic suppression occurring during the diapause maintenance phase (Wipking et al. 1995; Feder & Filchak 1999), with no subsequent significant independent effect of any other developmental phase. Thus, there will be a general 1:1 correspondence between individuals having greater metabolic suppression and later adult eclosion. A competing hypothesis is that the key factor dictating phenology is when diapause maintenance is terminated, regardless of metabolic depth, (Wadsworth et al. 2013) Alternatively, variation in adult emergence phenology may be driven solely by differences in the rate of post-diapause morphogenesis (Poseldovich et al. 2012; 2014). In each of these models, other components of diapause preparation, maintenance, and post-diapause development would be weakly correlated with eclosion time. Conversely, correlated shifts of different components within and between diapause phases can also lead to divergent phenology, especially if driven by directional selection. For example, earlier adult emergence timing may be driven by combinatorial effects of higher metabolic rate during the diapause maintenance phase, earlier termination of the diapause maintenance phase, and more rapid post-diapause development.

Here, we examine the role that these different components of diapause development play in the case of rapidly evolved phenological divergence in the apple maggot fly, *Rhagoletis pomonella* (Walsh), a text-book system for ecological speciation (Dres & Mallet 2002; Berlocher and Feder 2002). Divergent seasonal adaptation of the flies to synchronize adult emergence with their specific host fruits is well understood (Feder and Filchak 1999), making *R. pomonella* an excellent system for studying how different components of diapause development interact in the regulation of seasonal timing (Feder & Dombraski 2007; Ragland et al. 2017). Partially reproductively isolated populations that show consistent patterns of host-associated genetic differentiation, hereafter host races, are undergoing divergent selection to synchronize themselves with the fruiting times of their respective host plants. The derived host race shifted from infesting fruits of a native plant, the downy hawthorn (*Crataegus mollis*) to infest domesticated apples (*Malus pumila*) ∼160 years ago (Walsh 1861; Bush 1966). The apple varieties favored by *R. pomonella* ripen ∼1 month earlier in the season compared to hawthorns in the Midwestern US (Feder et al 1993, 1994). Apple and hawthorn flies spend most of the year in pupal diapause. Thus, the difference in adult and larval phenology between the races is determined by how flies transition from this shared state of pupal diapause during winter to completing adult development prior to emergence at different points the following summer and fall.

We investigate how variation in components of diapause maintenance and post-diapause development are associated with shifts in seasonal life history timing in the derived compared to the ancestral host race of *R. pomonella*. Specifically, we test whether the earlier phenology of the derived apple race is driven by alteration to individual components or multiple components of their diapause developmental trajectories. We use respirometric phenotyping to compare the post-winter metabolic rate trajectories of apple and hawthorn flies from sympatric populations under controlled laboratory common-garden conditions to identify the key phases of diapause development. Low metabolic rate is a reliable marker for the diapause maintenance phase that identifies individuals in diapause; metabolic rate then increases in a stereotypical trajectory during post-diapause development following the end of diapause (Ragland et al. 2009). We follow cohorts of flies from both host races in the laboratory from the cessation of simulated winter through to adult emergence, producing a time-series of metabolic rates for each individual. We fit these time series to mathematical models with parameters describing different aspects of diapause development corresponding with post-winter metabolic trajectories that include parameters modeling: 1) baseline metabolic rate during diapause maintenance, 2) timing of the transition from maintenance to post-diapause development, 3) metabolic rate at a plateau coinciding with early adult morphogenesis, 4) duration of the plateau, and 5) an exponential increase in metabolic rate corresponding with late adult morphogenesis culminating in adult emergence (Figure 1c). Comparisons across these individual models of metabolic trajectories allow us to test how patterns of post-winter diapause development differ between the two host races, and importantly test which component or combination of components is significantly associated with variation in adult emergence phenololgy. Thus, our approach provides a test of the hypothesis that variation in the duration of one or multiple components of the diapause-maintenance phase or post-diapause development account for earlier emergence timing in the derived apple host race. We also conduct a second experiment to specifically test whether overwintering and immediate post-winter diapause metabolic rates differ between the host races and how variation in these traits influences adult emergence timing.

**Figure 1.**
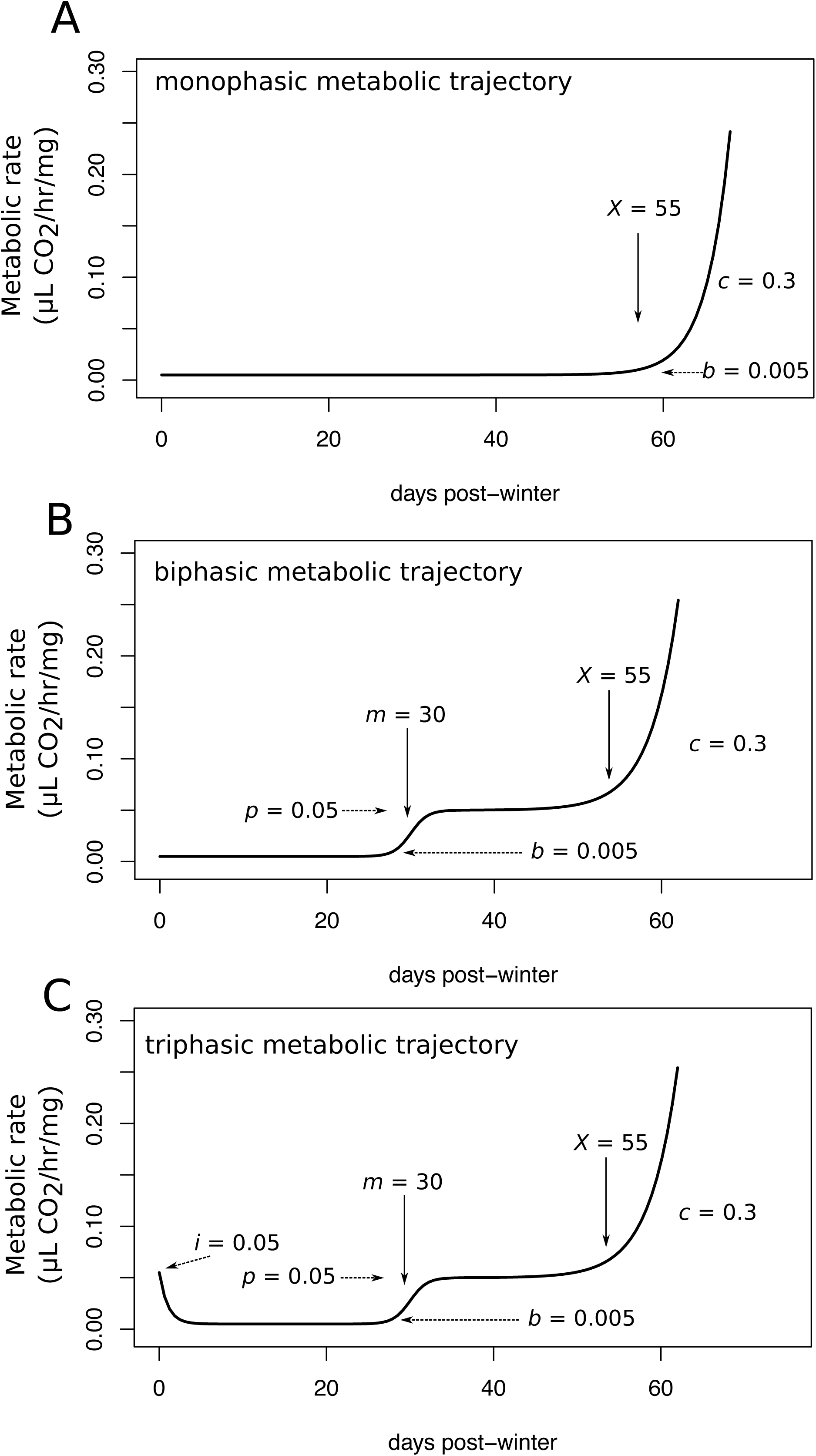
Examples of plotted metabolic trajectories following the A) monophasic, B) biphasic, and C) triphasic models described in the text. To demonstrate how different parameters affect aspects and the shape of metabolic trajectories and reflect different modules of diapause development, the values of the fitted parameters in each model are indicated in the plots with arrows pointing to their relevant position.

## Materials and Methods

### Insect collection and rearing

In the summer and fall of 2013 and 2014, apple and hawthorn race *R. pomonella* flies were collected from infested fruit from wild populations in Urbana, IL during peak infestation for both host plants. The apple and hawthorn sites are located ∼1,300 m apart, well within the flight range of these flies (Roitberg et al. 1984). Infested fruit were collected from the ground and trees and transported to the laboratory where they were kept in an environmentally controlled room under standard *Rhagoletis* rearing conditions (24°C and 14:10 Light:Dark (L:D): Filchak et al. 2000; Dambroski & Feder 2007). Fruit were placed in wire mesh baskets in plastic trays where wandering third instar larvae emerged naturally from the fruit and dropped into the trays. Trays were monitored daily for larvae, which were placed in petri dishes of moist vermiculite to pupariate. Petri dishes were kept at 24°C and 14:10 L:D for ten days. At the end of the ten days, pupae were transferred to clean petri dishes and placed in chambers with 85% humidity maintained by a saturated KCl solution (Winston and Bates 1960). Humidity chambers were then transferred to simulated winter conditions at 3.5° ±1.0 C, where they remained for 24 weeks. After removal from the cold, pupae were kept at 24° C, 14:10 L:D, and 85% humidity for the duration of the experiment.

### Measuring post-winter metabolic trajectories

Metabolic rates were measured using stop-flow respirometry (Lighton 2008), closely following the methods of Ragland et al. (2009). Upon removal from the cold, 96 pupae from each host race were weighed and placed into individual 5ml Norm-Ject^TM^ syringes (Air-Tite Products, Virginia Beach, VA, USA) fitted with three-way luer valves (Cole-Parmer, Vernon Hills, IL, USA). *Rhagoletis* pupae are small enough (∼5-11 mg) to reside within the top arm of the luer valve so that the syringe piston can be fully depressed. Syringes were flushed with CO_2_ free air, using an aquarium pump pushing air through a Dririte^TM^ (W.A. Hammond Co. Xenia, OH, USA) - Ascarite II^TM^ (Thomas Scientific, Philadelphia, PA, USA) - Dririte^TM^ scrubber column followed by an acidified (pH < 4.0) water bubbler to rehumidify the air. Syringes were sealed with 1ml scrubbed air, and the time of the syringe purge was recorded using Expedata logging software (Sable Systems International, Las Vegas, NV, USA). Four empty control syringes were also purged and sealed. Syringes were then returned to 25° C, 14:10 L:D incubators. After approximately 24 hours (exact time recorded for each individual), the full volume of the 1mL chamber was injected into a flow-through system with a Li-Cor 7000 infrared CO_2_ analyzer (Lincoln, NE, USA) interfaced with Expedata. The flow rate of the respirometry system was maintained at 150ml/min using an MFC-2 mass flow control unit (Sable Systems International) with a Sierra Instruments (Monterey, CA, USA) mass-flow valve, and the air in the system was scrubbed of CO_2_ and water using a Dririte^TM^ column followed by a Dririte^TM^-Ascarite II^TM^-Dririte^TM^ column. Water vapor from the injected sample was removed by an additional magnesium perchlorate column before entering the sample cell of the gas analyzer. Because both the purge and injection times were recorded in Expedata to the second, the exact time that each pupa spent sealed in the chamber was known and used in the subsequent analysis of respiration rates. Calibration of the gas analyzer was maintained during the experiment using pure nitrogen and a certified mixture of 465.8 ppm CO_2_ in nitrogen (Airgas, Radnor, PA, USA).

The respiration rate for each individual at each time point was calculated using the manual bolus integration method of Lighton (2008) with

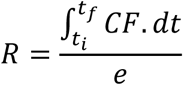

where *R* is the rate of respiration in μl CO2/h, *C* is the instantaneous concentration of CO2, *F* is the flow rate in *l*/*s*, *e* is the elapsed time interval in seconds between syringe purge and injection, and *t_i_-t_f_* is the time interval in seconds over which the injection bolus is detected by the gas analyzer. We conducted subsequent analyses on wet-mass equivalent metabolic rates in μl CO2/h/mg. Note that the expected variation in developmental state among individuals at a given time point precluded us from establishing a more complex allometric relationship between body mass and metabolic rate in this system.

### Analysis of metabolic trajectories

The previous study by Ragland et al. (2009) described a biphasic shape to *R. pomonella* post-winter metabolic trajectories, with a depressed baseline respiration rate during diapause punctuated by a logistic increase in CO_2_ production following the end of diapause that stabilizes at a plateau level before entering an exponential increase phase during later pharate adult development (Fig. 1b). Initial inspection of our data revealed that this basic biphasic pattern was present, but the majority of individuals in our study also revealed a characteristic increase in metabolic rate after transfer from cold to warm conditions, followed by an attenuation back down to a stable baseline (Fig. 1c). This initial increase in metabolic rate followed by a return to depression 2-3 days after, indicating pupae were still in the diapause maintenance phase, was also confirmed by an additional follow-up experiment described below. To account for this additional feature, we fit and compared three nested non-linear models to each individual fly’s time series of respiration rates: 1) a monophasic model consisting of an exponential increase from a single post-winter baseline metabolic rate (Fig. 1a):

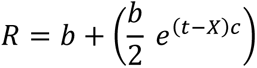

2) a biphasic model similar to the one described by Ragland et al. (2009), but redesigned so that the parameter values are more directly reflective of landmark time points and metabolic rates (Fig. 1b):

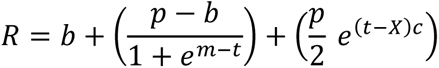

and 3) a triphasic model that also includes an initial post-winter elevated respiration rate followed by an immediate exponential decrease down to a baseline diapause rate (Fig. 1c):

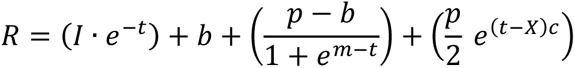

where *R* is the mass equivalent respiration rate at time *t, b* is the baseline post-winter metabolic rate, *p* is the plateau metabolic rate between diapause termination and pharate adult development, *X* represents the timing of the exponential increase in respiration during pharate adult development (equal to the time at which the metabolic rate is 2x higher than the previous stable rate *b* or *p*), *m* is the timing of the logistic increase during diapause termination, *I* is the initial elevated post-winter metabolic rate, and *c* is a scaling factor for the slope of the exponential phase. Initial AIC analysis during model development did not support the inclusion of additional scaling parameters for the logistic increase or initial decrease phase for any individuals.

All three models were fit for each individual in R 3.0.1 (R Development Core, Vienna, Austria), using the *nls* function. Starting values for each parameter were obtained from visual inspection of the metabolic trajectories; each parameter except for c models either a metabolic rate or time, and thus a position on the y or x axis of the metabolic trajectory plot, respectively (Fig. 1). Parameters were constrained to being biologically reasonable (e.g., all time and metabolic rate parameters > 0). Models were compared using AICc in the R package MuMIN (Barton 2015). Goodness of fit of the models selected by AICc was assessed by examining fitted curves and the distribution of residuals.

### Testing whether variation in adult emergence timing is associated with variation in the same developmental trajectory model parameters across both host races

If particular phases of diapause or post-diapause development account for differences in adult emergence phenology, then inter-individual variation in parameters from the above models representing these phases should associate with adult emergence timing 1) within, and 2) between fly host races. Due to strong covariance among diapause trajectory parameters, as well as collinearity between some parameter estimates and their errors (as expected in this case where the degrees of freedom in each trajectory model vary exactly with phenology because flies that take longer to emerge are measured more times), we relied primarily on a partial correlation approach rather than multiple linear regression for statistical testing. We conducted partial correlation analyses assessing the relationships of the parameters *b*, *m*, *p*, *X-m*, *c* to days to adult emergence using the R package ppcor (Kim 2015). Analysis of variation was conducted both across the host races considering host as a binary variable, and within apple and hawthorn flies considered separately.

### Analysis of over-wintering and immediate post-winter metabolic rates

The unexpected presence of elevated metabolic rates in the first few post-winter measurements 2-3 days after removal from cold combined with early diapause termination of many apple flies introduced more variability into the initial post-winter baseline measurements than expected in the original experiment. To better control for the rapid change in metabolic rates during this immediate post-winter phase and to further examine the dynamics of the immediate post-winter metabolic increase and subsequent decline, we performed a second metabolic phenotyping experiment the following year. Here, we measured metabolic rates of 40 apple race and 40 hawthorn race pupae from the same populations at Urbana, Illinois (but collected during the 2014 field season). Larvae and pupae were handled as above except that we used a 28-week simulated winter. At the end of this 28-week cold period, we placed the pupae in syringes, purged the syringes with CO_2_ free air as described above and sealed them under continuous simulated winter conditions (3.5 °C) for 24 hours. After 24 hours, syringes were placed in an insulated cooler with ice packs and brought to our room-temperature respirometry system. Sealed syringes were kept in the cooler until immediately before injection, resulting in a measurement of CO_2_ produced by each individual under uninterrupted winter temperatures. The post-winter metabolic rate of the flies was then measured for 12 days at 24°C as described above with the exception that measurements were made every 24 hours. Syringes were re-purged immediately after injection. Metabolic rates were calculated as described above. Pupae were then kept in humidity chambers in 0.2 ml perforated tubes and monitored daily for adult emergence.

### Statistical analyses of immediate post-winter metabolic rates

Our analyses of this second experiment had two distinct goals: 1) to determine whether the two host races differ in their metabolic profiles over this dynamic period of immediate post-winter metabolic change, and 2) to determine the extent to which metabolic rates during the overwintering and immediate post-winter phase may influence the timing of adult emergence. To test whether and at which time points the host races differ in their immediate post-winter diapause metabolism, we conducted a repeated-measures ANOVA of daily metabolic rates for each fly as a function of host race to determine metabolic time points that differed significantly between apple and hawthorn flies. Based on those results, we then asked how variation in two key post-winter time points that differentiated the host races, specifically 1 day and 10 days out of overwintering, and variation in the cold overwintering metabolic rates covaried with each other and with adult emergence phenology using a partial correlation analyses, as above.

## Results

### Metabolic rate trajectories suggest stereotypical developmental trajectories

Apple host race flies emerged as adults significantly earlier than hawthorn host race flies with mean and median eclosion times for the apple race flies at 51d and 52d (SE = 2.33, IQR = 38.25 – 56.75), respectively compared to 81d and 79d (SE = 2.98, IQR = 73 - 87.5) for the hawthorn race (Fig. 2a, KS test, D = 0.793, P < 0.00001). The majority of pupae followed either a biphasic or triphasic metabolic rate trajectory model based on AICc model selection (Fig. 2b). However, there was a single fly showing the monophasic trajectory that emerged only 19 days after simulated winter. AICc results for each fly are presented in Table S1, and the best-fit models for each fly are presented in a file in the online supplementary material and Figure S1. Apple flies had more metabolic rate trajectories following the biphasic model fit, whereas, hawthorn flies had a roughly equal ratio of bi- and triphasic model fits (G-test, G_(2*df*, N = 68)_ = 16.962, p < 0.001), Table S1). The lower frequency of triphasic trajectories in apple flies was likely driven by their shorter time to adult emergence. With fewer data points during the baseline metabolic rate phase of the trajectories, apple flies tended to have less power to fit the initial decrease curve, with the logistic increase of diapause termination often beginning before the decrease phase stabilized (Fig. S1).

**Figure 2.**
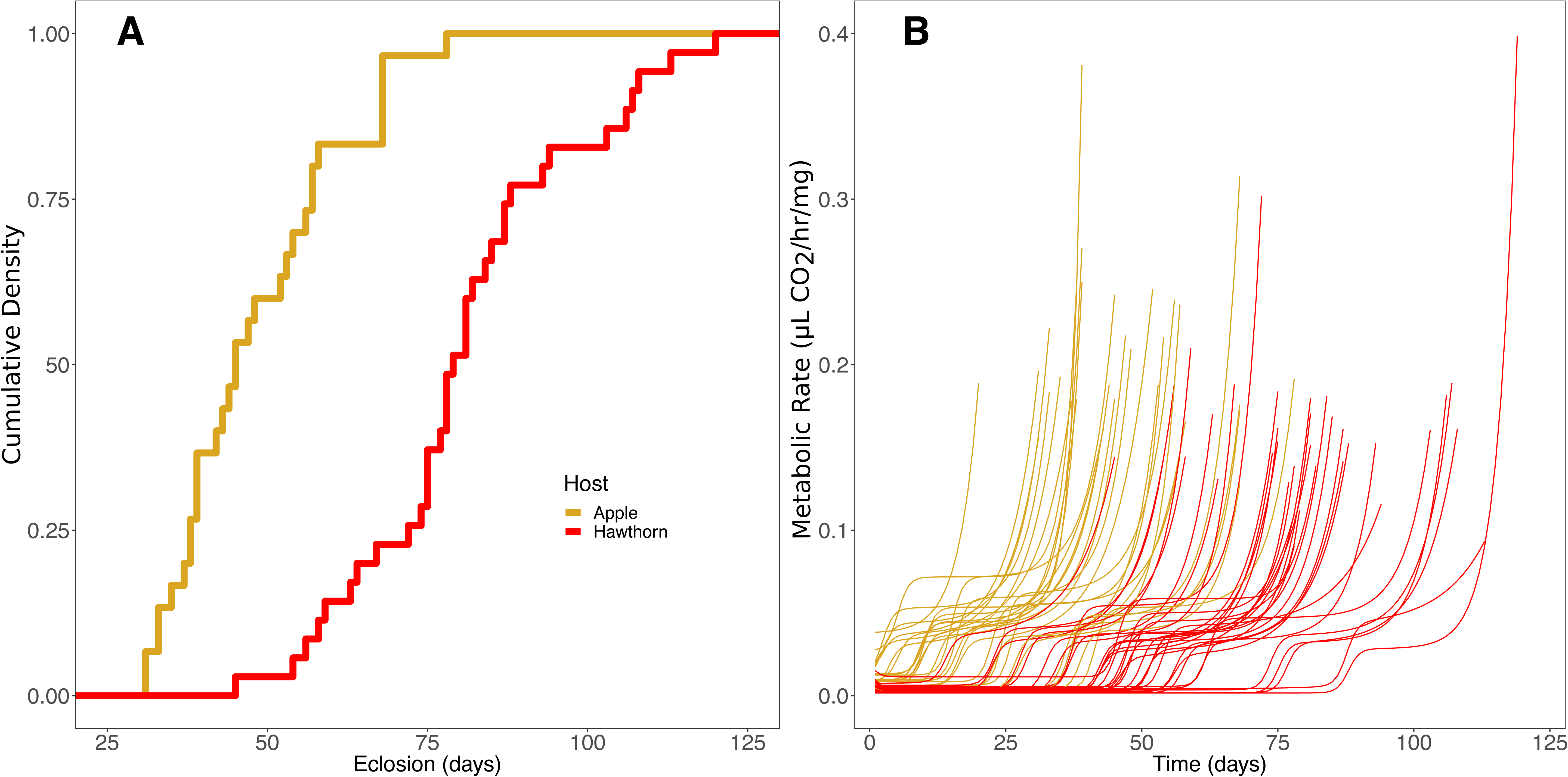
Plots showing A) that apple flies (yellow) emerge as adults earlier than hawthorn flies (red), and B) the best-fit non-linear models of the metabolic trajectories for each apple fly (yellow) and hawthorn fly (red) that survived to adult emergence in experiment 1.

The pattern of initial decrease of metabolic rates from an immediate post-winter peak was present in both hawthorn and apple flies (Fig. S1 and see results of second experiment below). Curves from the fitted models (Fig. 2b) generally fit the shape of the data closely (Fig. S1). The mean residual standard errors *S* were low for the best-fit models, with a mean of 0.0041 μl CO2/h across all flies and a range of 0.0012 – 0.0089 μl CO2/h. We note that R^2^ is generally not an appropriate metric of goodness-of-fit for non-linear regressions (Spiess & Neumayer 2010). To use parameter estimates from the best fit models for each fly in subsequent analyses, we tested whether the subset of nested shared parameter estimates differed between the hierarchical biphasic and triphasic models fit for the same individuals. These shared parameters: baseline metabolic rate during the diapause maintenance module (*b*), timing of the end of the diapause maintenance module (*m*), post-diapause plateau metabolism (*p*), post-diapause plateau duration (*X*), and exponential scaling (*c*) did not differ between the bi- and tri-phasic models in paired t-tests (Table S2).

### Apple and hawthorn flies differ in key post-winter metabolic trajectory parameters

While the host races share the same basic metabolic trajectory templates, apple and hawthorn flies differed in parameters related to both phase-specific metabolic rates and, most notably, the timing of the transition out of the diapause maintenance phase. Estimates of three parameters: baseline metabolic rate *b*, the timing of the transition out of the diapause maintenance phase *m*, and the post-diapause plateau metabolic rate *p*, differed between host races (Fig. 3a-c; two-tailed t-tests, t = 4.0787, P = 0.0026; t = −8.61, P = 3.64×10^-12^; t = 3.8012 P = 0.000385, respectively). Neither the relative timing of exponential metabolic increase during adult development X-m nor the exponential scaling parameter c were significantly different between host races (Fig. 3d-e; t-tests, t = −0.39081, P = 0.6973; t = 1.444, P =0.1542, respectively). Compared to hawthorn flies, apple flies had higher baseline metabolic rates, earlier diapause termination times, and higher post-diapause metabolic plateaus (Fig. 3a-c).

**Figure 3.**
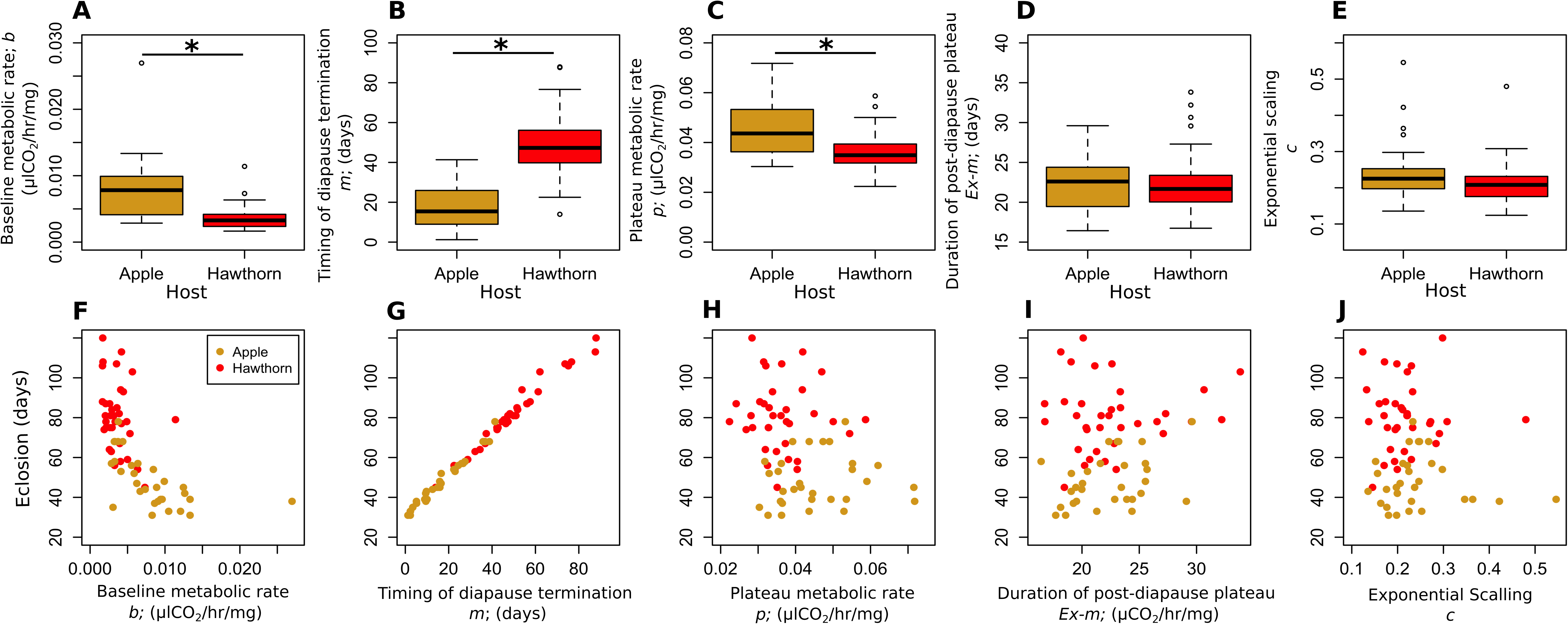
Box plots comparing fitted parameter values that show: A) apple flies have a higher baseline metabolic rate (*b*) than hawthorn flies, B) apple flies have earlier timing of diapause termination (*m*) than hawthorn flies, C) apple flies have higher post-diapause plateau metabolic rates (*p*) than hawthorn flies, D) apple and hawthorn flies do not differ in post-diapause plateau duration (*X-m*) nor E) exponential scaling (*c*). The box plots are followed by scatter plots of the relationships between adult emergence time on the y-axis and F) baseline metabolic rate (*b*), G) timing of diapause termination (*m*), H) post-diapause plateau metabolism (*p*), I) post-diapause plateau duration (*X-m*), and J) exponential scaling (*c*) with individual apple flies indicated in yellow and hawthorn flies in red, showing that adult emergence timing is strongly correlated with the timing of diapause termination (*r* = 0.994) with much weaker relationships to the other fitted metabolic trajectory parameters.

### Associations between model parameters and emergence timing within host races

Our correlation analysis revealed a strong relationship across both host races between *m*, the timing of the end of the diapause module, and adult emergence timing across both host races (*r* = 0.994, P = 1.07e^-57^, Fig. 3G, Fig. 4). Earlier timing of the end of the diapause maintenance module was tightly correlated with earlier emergence timing (Fig. 3G). Weaker but significant partial correlations with adult emergence timing also existed between post-diapause plateau metabolism (*p*), metabolic rate duration of the post-diapause plateau (*X*-*m*) and the slope of the exponential increase phase *c* (*r* = −0.37; P = 0.0013; *r* = 0.772, P = 5e^-13^, and *r* = −0.533, P = 0.00015, respectively; Fig. 3I-J; Fig. 4). The baseline metabolic rate *b* did not show significant direct correlations with adult emergence time (Fig 3F; Fig 4). These patterns were consistent both within and across the two host races (Fig 3F-J; Fig. 4; Fig S2). The strength of the correlation between *m* and adult emergence timing was nearly identical when the partial correlation analyses were run separately for each host race (apple *r* = 0.98 and hawthorn *r* = 0.99; Fig S2), reinforcing the importance of the timing of the end of the diapause maintenance module as the primary driver of phenological divergence between the apple and hawthorn host races.

**Figure 4.**
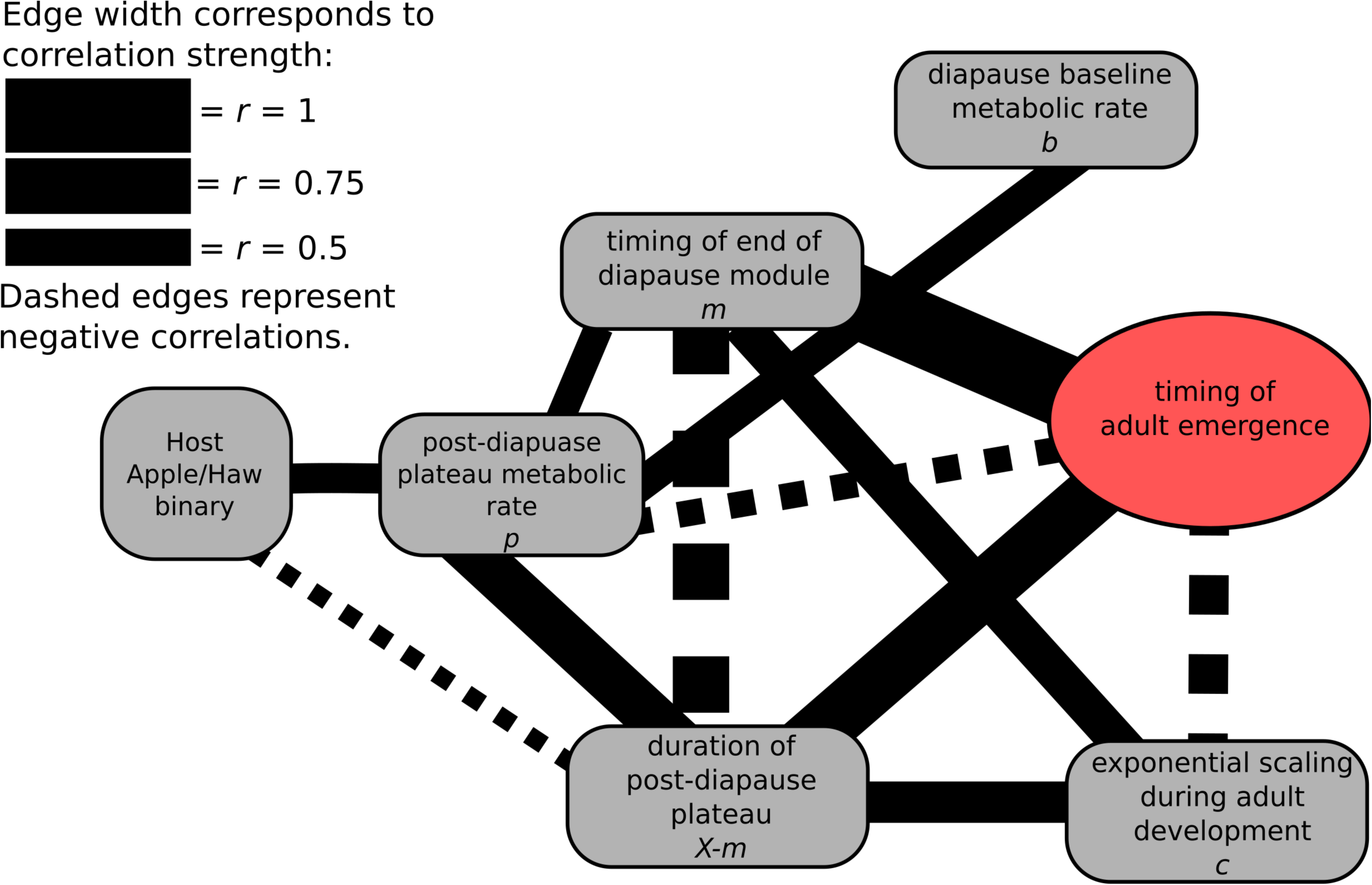
Partial correlation network showing the major contribution of the timing of diapause termination (*m*) *(r* = 0.994) along with the minor contribution of post-diapause parameters describing the plateau metabolic rate (*p*), the duration of the post-diapause plateau (*X-m*), and scaling of the exponential phase (*c*) (*r* = −0.37, *r* = 0.772, and *r* = 0.533, respectively) with adult emergence timing as well as additions significant partial correlations among parameters. Edges depict all significant (P < 0.05) partial correlations. Edge weights are proportional to the magnitude of Pearson’s *r*, for both positive correlations (solid lines) and negative correlations (dashed lines).

### Host races diverge in how they respond to initial favorable conditions

The overall shape of the metabolic trajectories for both host races in the second respirometry experiment was consistent with the dynamics found in our first experiment. The end of simulated winter and transfer to 25 °C resulted in a sharp initial metabolic increase that peaked at 24-72 hours after removal from the cold and subsequently declined to a baseline metabolic rate over the course of 10 days (Fig. 5). Overwintering metabolic rates (measured in the cold) were nearly an order of magnitude lower than post-winter baseline metabolic rates from the experiment above, mean 0.00115 vs. 0.01035 μlCO2/hr/mg, respectively, and did not differ between the host races (Fig. S3). However, the two host races did differ in their immediate response to warming, with apple flies being more metabolically responsive to warming than hawthorn flies (repeated measures ANOVA: time x host, F_12,552_ = 3.54 p <0.001; Table S3; Fig. 5). Thus, apple flies were more metabolically responsive to the shift from cool simulated over-wintering conditions to warm post-winter conditions, rising to higher metabolic rates than hawthorn flies after simulated overwintering, particularly on days 1 & 2 post-winter, with some individuals appearing to terminate diapause before day10 (Fig. 5).

**Figure 5.**
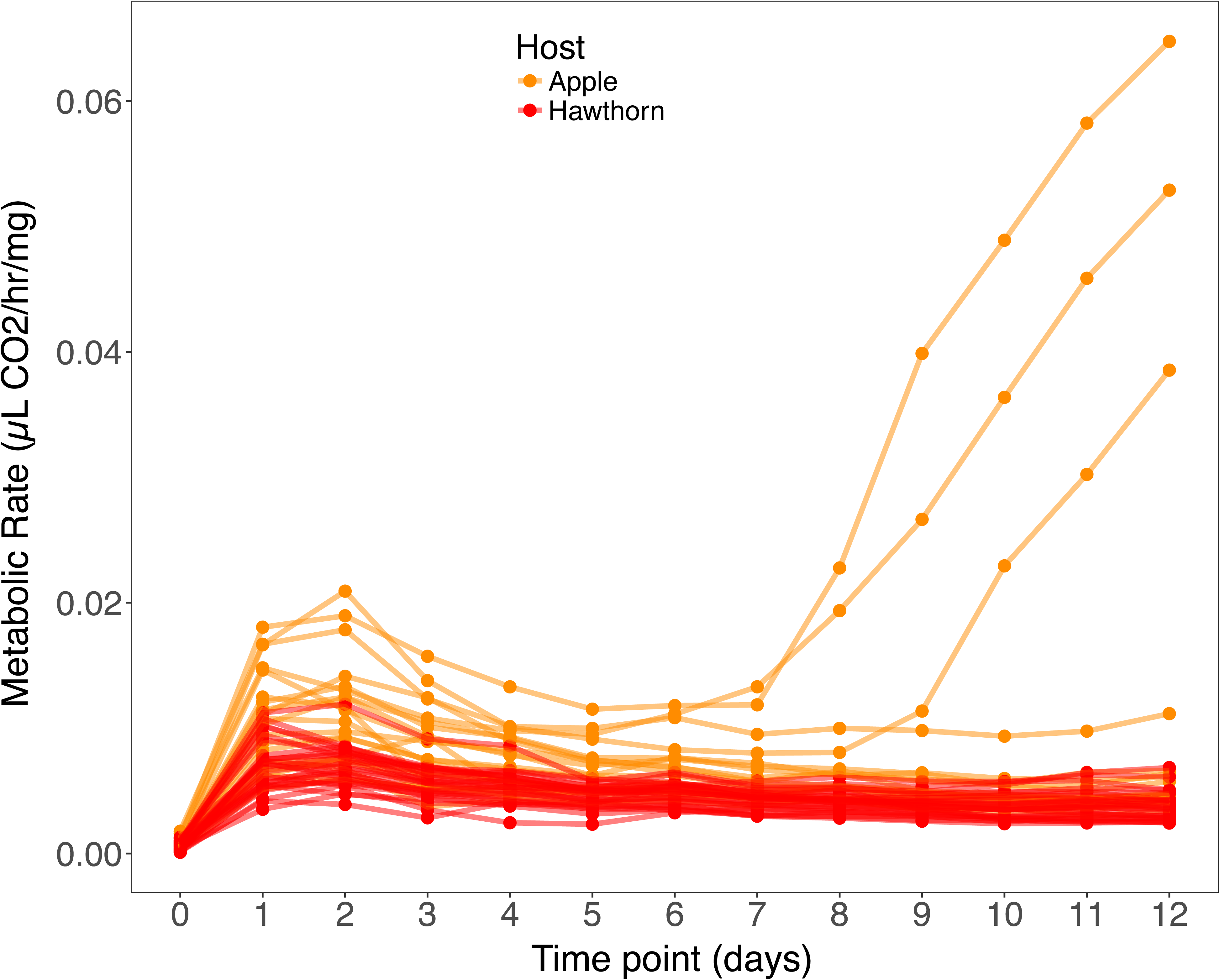
Plots of metabolic rate every 24 hours over 12 days for surviving apple flies (yellow) and hawthorn flies (red) from experiment 2 showing the immediate post-winter metabolic increase over days 1 and 2 exhibited in both host races, with apple fly pupae increasing their metabolic rates substantially higher than hawthorn fly pupae after removal from the cold. After this initial metabolic increase, all hawthorn pupae and most apple pupae enter back into metabolic suppression at baseline post-winter diapause maintenance levels. Time zero on the x-axis represents the overwintering metabolic rate measurement over which flies were purged and measured under continuous simulated winter conditions. See Fig. S4 for a box plot depicting variation in overwintering metabolic rates in more detail.

Partial correlation analysis of overwinter, day 1, and day 10 metabolic rates with adult emergence time in the second experiment showed significant but weaker overall covariance among variables than was found in the first experiment (Fig 6). Overwintering metabolic rate was weakly but significantly positively correlated with metabolic rates at the two other time points day 1 and day 10 after removal from cold, as well as host (*r* = 0.37, P = 0.008, *r* = 0.346, P = 0.013; *r* -= 0.315, P = 0.026, respectively, Figs. 6, S4). The only variable with a direct correlation to adult emergence timing was an inverse relationship with day 1 metabolic rate (*r* = −0.318, P = 0.025, Figs 6, S4). Pupae of both host races that were more metabolically responsive during the rapid rise in metabolism after shifting from cool overwintering conditions to warm post-winter conditions were more likely to emerge earlier as adults. This correlation was much weaker than that observed between adult emergence timing and the timing of the end of the diapause maintenance module (*r* = −0.318 vs *r* = 0.994), indicating that if day 1 metabolic rate influences adult emergence phenology, it likely does so through its effect on the timing of the end of diapause maintenance.

**Figure 6.**
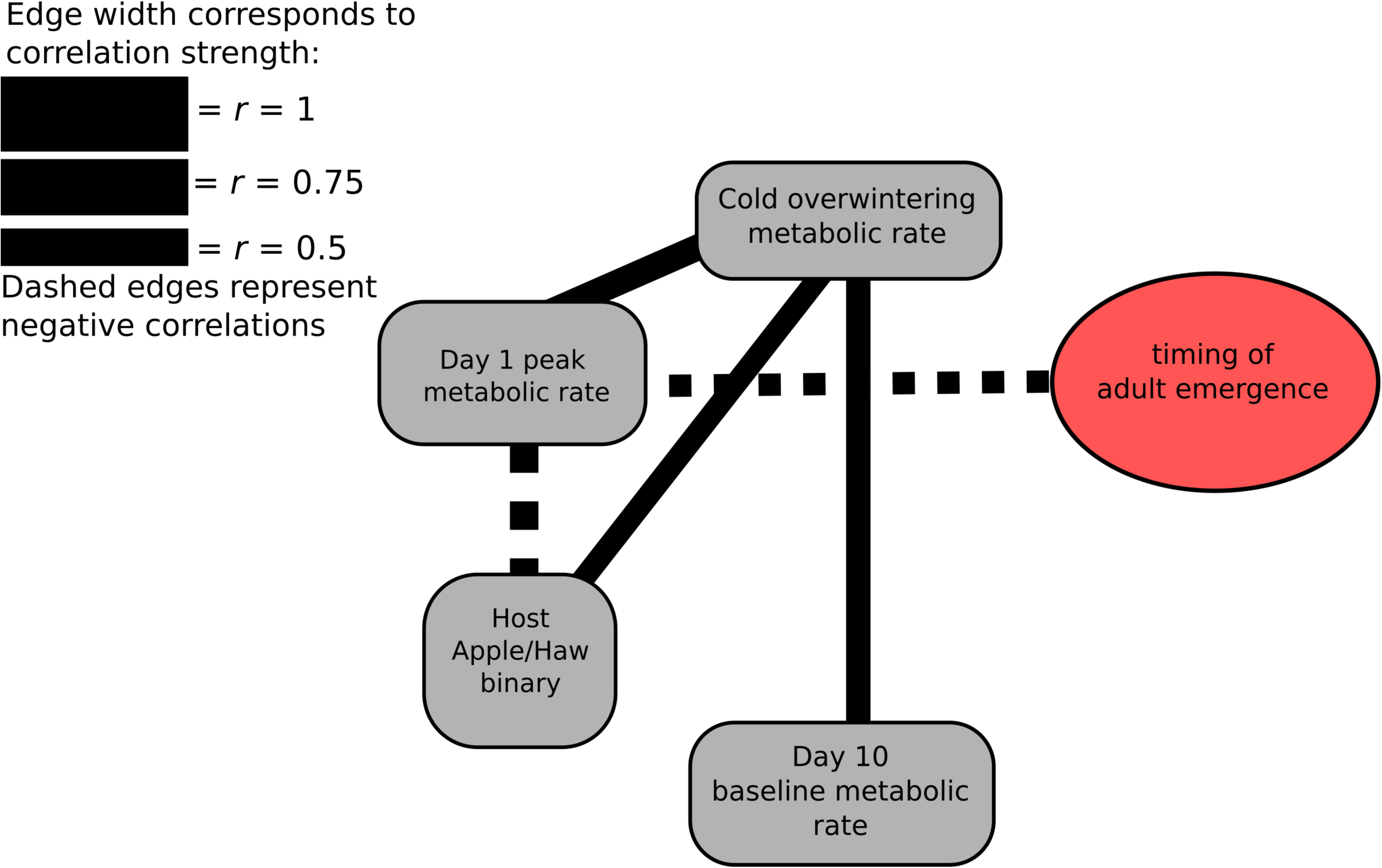
Partial correlation network showing the relatively weak negative correlation between day 1 metabolic rate and adult emergence timing (r = −0.318), as well as additional significant partial correlations among overwintering metabolic rate and post-winter metabolic variables in experiment 2. The three metabolic rate variables over this period are modestly positively correlated, and pupae with particularly strong metabolic increases at day 1 tended to emerge earlier as adults. Edges depict all significant (P < 0.05) partial correlations among variables. Line weights are proportional to the magnitude of Pearson’s *r*, for both positive correlations (solid lines) and negative correlations (dashed lines).

## DISCUSSION

Diapause is an important component of adaptation to temperate environments for most insect species, and it may often serve as a critical step governing the seasonal phenological timing of insect life histories (Tauber & Tauber 1981). As a multi-faceted, alternative developmental pathway, diapause is not just a static resting state. Diapause is a complex trait, possessing a number of physiologically distinct functional components that are expressed along the course of diapause development (Kostal 2006). Thus, the ability for insect populations to rapidly evolve new phenologies is likely dependent on the complexity with which these different components of diapause development interact to affect overall seasonal timing. Here, we show that although apple and hawthorn flies differ in a number of timing and metabolic rate parameters of post-winter development (Figs 3A-E), variation in adult emergence, both between and within the host races, is primarily due to differences in the timing of the transition between diapause maintenance and post-diapause development (parameter *m*; Fig 3F-J; Fig 4). In both host races, adult flies emerged on average 31.6 days after diapause termination, with a standard deviation of 2.4 days. Wadsworth et al. (2013) reported similar results for a primary effect of diapause termination on adult emergence time in the univoltine-Z and bivoltine-E strains of the European corn borer, another system where divergence in seasonal phenology drives allochronic isolation (Dopman et al. 2010), perhaps highlighting the general importance of diapause termination timing in the evolution of insect phenology.

In addition to diapause termination, adult emergence timing in the host races is related, although to a much lesser degree, to two other post-winter development parameters: 1) the duration of the post-diapause plateau (*X-m*) when early pharate adult morphogenesis is occurring (Ragland et al. 2009); and 2) the slope of the exponential increase phase of pharate adult development (*c*) late in the process of adult morphogenesis (Figs 3H-J). In our second experiment, a weak but significant inverse correlation was also observed between adult emergence and the initial day 1 post-winter metabolic rate, such that pupae that more strongly increase metabolic rates after the transition from cool over-wintering temperatures to warm post-winter temperatures are more likely to emerge as adults earlier (Fig 6). However, the timing of the end of the diapause maintenance phase appears to be the major factor driving variation in overall adult emergence phenology, with the later-acting post-diapause development parameters having much subtler effects on any residuals variation in emergence timing (Figs 3H-J, Fig. 4).

The overall covariance structure presented in the partial correlation network (Fig. 4) indicates that the major effect of the timing of the transition between the diapause maintenance and post-diapause development phases on phenology is not entirely independent of the other metabolic trajectory parameters. Although the timing of transition between phases appears to be the primary driver of phenology, this variation may not act in a singularly modular fashion, as we would infer if the parameters correlated with adult emergence timing did not otherwise overlap in the partial correlation network. However, most of the correlations among metabolic trajectory parameters were relatively modest (only two were stronger than *r* = 0.5, with the strongest being between *m* and *X-m* at *r* = −0.76), implying that their respective responses to selection may not be highly constrained. Thus, the covariance between diapause termination and post-diapause development is not absolute, both the diapause maintenance phase and the post-diapause development phase may be capable of separate responses to selection on adult life history timing, a result that has broader implications for phenological adaptation in many contexts from seasonal synchronization with novel hosts to rapid adaptation to new climatic regimes during introductions or range expansion to shifting seasonality as a result of climate change (e.g. Bradshaw & Holzapfel 2001; Ording & Scriber 2005; Gomi et al. 2007; Wadsworth et al. 2013).

In certain insects, variation in diapause phenotype may have relatively simple genetic architecture. Diapause-mediated allochronic isolation in the European corn borer is under the control of a segregating locus of major effect on the Z-chromosome (Glover et al. 1992; Dopman et al. 2005; Wadsworth et al. 2015). Similarly, variation in diapause induction appears to be driven largely by a large-effect Z-linked locus in a butterfly species complex comprised of *Paplio glaucus*, *P. canadensis*, and *P. appalacheinsis* (Hagen & Scriber 1989; Kunte et al. 2011). But latitudinal variation in life history timing in *Papilio* butterflies may also be associated with autosomal regions as well (Ryan et al. 2017). In contrast, geographic variation in photoperiodic sensitivity for diapause induction in the mosquito *Wyeomia smithii* appears to be under more complex genetic control, involving multiple loci of moderate effect with strong epistasis (Lair et al. 1997; Mathias et al. 2007). Genetic variation in life history timing in *R. pomonella* appears to be highly quantitative, showing widespread statistical associations with markers across the genome (Filchak et al. 2000; Michel et al. 2010; Egan et al. 2015; Ragland et al. in 2017). Our results suggest that the many genes implied to affect adult emergence phenology primarily do so by their effect on the timing of termination of the diapause maintenance module. Thus, timing of diapause termination is a developmental hub through which quantitative allelic variation flows to modify adult emergence phenology.

The genetic complexity of life history timing in *R. pomonella* might seem at odds with the relative simplicity of diapause termination primarily dictating adult emergence timing. However, there is good reason to suspect that many genes or gene networks may play a role in driving variation in the duration of the diapause maintenance module. The pre-winter initiation of diapause is not simply a switch ceasing development, and the termination of diapause is not simply a switch resuming development. Rather the phases are each characterized by a progression of physiological changes that lead up to each transition along the developmental pathway (Denlinger 2002; Kostal 2006; Kostal et al. 2017). The dynamic nature of this developmental trajectory allows for many different transitions at which allelic variation might influence diapause timing, and many intermediary developmental avenues exist along which highly polygenic variation could act additively to affect eclosion timing. Precisely how each of these processes and events relate to the detectable metabolic increase during diapause termination is not currently known. Further characterizing variation in these intermediate mechanistic steps to diapause termination will help parse the complex pattern of genomic variation associated with life history timing in these flies.

The earlier diapause termination of apple than hawthorn flies may partially be due to differences in their developmental progression during the overwintering portion of the diapause maintenance phase. Apple pupae may become more primed or potentiated to respond to environmental cues sooner after winter than hawthorn pupae. For example, apple flies may be prone to enter or progress further into the stage of diapause development permissive for termination of the maintenance phase earlier in the winter compared to hawthorn flies. Alternatively, the host races could emerge from winter in the same state of diapause development, with the difference in adult emergence time resulting from variation in their respective responses to subsequent environmental cues. Although the two host races did not show a detectable difference in overwintering metabolic rate, the fact that apple flies were more metabolically responsive to the shift from cool overwintering temperatures to warm post-winter temperatures, and the very rapid nature of diapause termination in many of the apple flies, argues that apple fly pupae are potentiated to respond to diapause-termination cues earlier than hawthorn fly pupae. The finding of higher initial metabolic response to warming in apple flies combined with the association between earlier adult emergence and higher initial metabolic rates just after warming in both host races suggests a modification to the simple hypothesis that basal metabolic rates during diapause themselves are a primary driver of diapause duration (Wipking et al. 1995; Feder & Filchak 1999). Instead, perhaps the dynamic responsiveness of metabolism to thermal shifts indicates greater potentiation to respond to diapause termination cues both within and between the host races and acts upstream of the timing of the end of diapause maintenance.

A recent RNAseq study comparing the transcriptomes of the host races during simulated overwintering and soon after removal from the cold is consistent with the hypothesis that apple flies are more potentiated to terminate diapause after winter than hawthorn flies (Meyers et al. 2016). The head transciptomes (including neuro-endocrine tissues) of apple and hawthorn flies were found to already be strongly differentiated before pupae were removed from the cold, with biases towards up-regulation of genes involved in cell proliferation and development in the apple flies (Meyers et al. 2016). Our finding of greater responsiveness of metabolic rate in apple than hawthorn race pupae as they shift from cold, developmentally suppressive temperatures to warm, developmentally permissive temperatures also suggests apple race pupae become more potentiated for diapause termination by the end of simulated winter treatments compared to hawthorn race pupae.

The immediate post-winter metabolic increase and subsequent decline observed in our second experiment (Fig. 5) is a novel finding with implications for how dormancy is maintained in the face of abiotic conditions that are otherwise permissive for non-diapause development. Diapause has been most extensively studied in insects with multivoltine life cycles (e.g. Schmidt et al. 2008; Ragland et al 2010; Emerson et al. 2010; Bao et al. 2011; Lehmann et al. 2014). For multivoltine organisms, trophic resources tend to have broad temporal distributions, and selection favors fast resumption of non-diapause development as soon as abiotic conditions are physiologically permissive. Thus, it is not surprising that for multivoltine species, the diapause maintenance phase typically terminates during winter, with insects remaining in post-diapause quiescence (dormant due to physiologically restrictive abiotic conditions rather than programmed metabolic and developmental suppression) until physiological temperature thresholds are surpassed, after which time they initiate post-diapause development (Hayward et al. 2005). In contrast, univoltine insects with narrow resource windows like *R. pomonella* derive no benefit from terminating diapause maintenance early during the winter because they must remain dormant even after conditions become permissive the following spring until it is time to initiate the post-diapause developmental phase to synchronize themselves with host fruit phenology.

The response of *R. pomonella* to the end of simulated winter was therefore more metabolically dynamic than we expected (Fig. 5). The shift from overwinter respiration to post-winter diapause respiration was neither a gradual linear increase nor a discrete stair-step up to the post-winter level. Rather, pupae showed a rapid but transient metabolic increase immediately after removal from the cold, peaking around 48 hours at a level 4 to 5 fold higher than the eventual post-winter baseline metabolic rates (Fig. 5). During normal development, metabolic rates of ectotherms increase with temperature (Angiletta 2009). Thus, thermally sensitive metabolic rates can lead to increased energy consumption during warmer conditions, which may prove costly to univoltine insects that need to conserve energy stores for prolonged dormancy (Irwin & Lee 2000; Mercader & Scriber 2004; Kostal et al. 2011; Vrba et al. 2014). However, diapausing insects may be able to mitigate these energetic costs by reducing the thermal sensitivity of their metabolic rates (Williams et al. 2012; 2015), as we saw in our trials when initially high metabolic rates were reduced within days of transfer to warm, permissive thermal conditions. Active suppression of metabolism in the face of physiologically permissive temperatures may play an important role in maintaining optimal overwintering energetics in the face of intermittent periods of warmth both in the fall and spring (Hahn & Denlinger 2011; Sinclair 2015). Differences in the immediate post-winter metabolic dynamics of apple and hawthorn flies might therefore represent an additional axis of divergent seasonal adaptation in this classic speciation system that may also be important to disentangle in other systems with rapid adaptation to new phenological regimes.

## CONCLUSIONS

Specialist insects make up a large proportion of metazoan biodiversity (Jaenike 1990; Bush & Butlin 2004). How insect life history timing evolves has important implications for both the origin and maintenance of a large swathe of biodiversity (Bush 1993). The tempo of seasonal adaptation for many species likely depends on the covariance structure of different components of dormancy regulation and how these components act to influence phenology. Are different facets of diapause development free to evolve independently, or are shifts in life history timing constrained by a need for simultaneous evolution across several phases of diapause development? Here, we found that termination of the diapause maintenance phase plays a particularly important (but not sole) role in synchronizing insect phenology with that of ephemeral resources (Figs 3,4). In the case of *R. pomonella*, complex patterns of quantitative allelic variation appear to act primarily through the hub of diapause termination to affect seasonal timing, with additional input via post-diapause development. The dual roles of these two developmental phases may be a coarse and fine adjustment of phenology, respectively. It is important to note that the magnitude of effects of diapause termination and post-diapause development reported here were measured under a simplified laboratory temperature regime: a constant post-winter temperature of 25 °C. It remains to be determined whether the effects of either of these phases on eclosion phenology are themselves function-valued traits driven by temperature. One possibility is that the duration of the diapause maintenance phase sets life history timing broadly across years and that the post-diapause development module may allow fine-tuning of adult emergence timing to within year variation in host-plant phenology. For example, apples always fruit earlier than hawthorns in our sympatric field sites. However, in some warm years, apples and hawthorns both fruit earlier and in cooler years both fruit later. Metabolic rates during the post-diapause developmental phase are substantially higher than during the diapause maintenance phase and we speculate that post-diapause development may be more responsive to temperature acting to fine tune synchrony with host plant availability and local abiotic conditions from year to year at any particular location and across a latitudinal gradient of seasonal conditions throughout this fly’s range. Future work on the thermal sensitivity of specific modular components across these two modules of the diapause developmental trajectory is needed to test this hypothesis for synchronizing adult emergence timing with weather-related local variation in host fruit phenology.

In *Rhagoletis* flies, variation in seasonal adult emergence timing is associated with on-going divergent adaptation to the novel apple host plant and reproductive isolation, but this shift to earlier seasonal timing is analogous to expectations for phenological shifts under climate change in coming decades. A primary motivation of this research was to determine the extent to which the inherent complexity of diapause development is likely to aid or constrain the rapid evolution of seasonality in univoltine insects. In the case of these two host races of *R. pomonella*, the partially independent major and minor effects related to the timing of the transition out of diapause maintenance and components of post-diapause development bodes well for the continued evolvability of adult eclosion time in this system. If phenological adaptation in these flies is not primarily constrained by strong antagonistic pleiotropy among different facets of diapause development, it may be limited more by available genetic variation affecting each intermediate phenotype. However, the apple and hawthorn fly speciation story is superimposed on strong local phenological adaptation of these flies along latitudinal clines as well (Dambroski & Feder 2007; Michel et al. 2010; Doellman et al. 2018a, b), which may act as a vast reservoir for standing genetic variation in seasonality traits across the continent (Powell et al. 2013). The combination of relatively simple, major effect basis for variation in adult emergence phenology coupled with abundant genetic variation may mean that natural selection has a fairly free hand with these insects to keep pace with changing seasonal conditions or colonize novel niches. However, the existence of moderate antagonistic covariance among traits with links to phenology may potentially constrain the evolution of novel phenotypes outside the current trait distribution. Furthermore, elucidating the roles of diapause maintenance and post-diapause development as important intermediate phenotypes driving differences in phenology sets the stage for future studies to begin drawing stronger connections between allelic variation across the genome and the complex ecological traits of life history timing, local adaptation, and ecological reproductive isolation. Given that genetic variation in the diapause maintenance phase (Wadsworth et al. 2013) or in post-diapause development (Poseldovich et al. 2012; 2014; Stalhansske et al. 2014) are thought to be independent drivers of adaptive shifts in phenology in other insect systems, we expect that the dual role for diapause and post-diapause development in regulating life history timing may be partially modular across a wide range of insect species. Thus, the idea that phenologies may be tuned to local seasonality by a combination of genetic and environmental effects (GxE) is broadly applicable across diverse groups of insects and germane to understanding persistence versus declines of insect populations in the future as seasonality is further altered by anthropogenic change.

## Acknowledgements

We would like to thank Meredith Doellman, Glen Hood, and Pete Meyers for help with field collections. Caroline Williams provided invaluable advice on our respirometry configuration. Help with phenotyping was provided by Jennifer Serviss, Caelen Schreiber, Andre Szejner, Chao Chen, Genevieve Comeau, and Xiaoping Wang. This research was supported by NSF IOS 170773 to GJR and JLF, NSF DEB 1638997 to GJR, NSF DEB 1638951 to JLF, and NSF DEB 1639005 to THQP and DAH, as well as NSF IOS 1257298, the Florida Agricultural Experiment Station, and the joint FAO/IAEA CRP Dormancy Management to Enable Mass-rearing to DAH.

---

**Figure S1.**
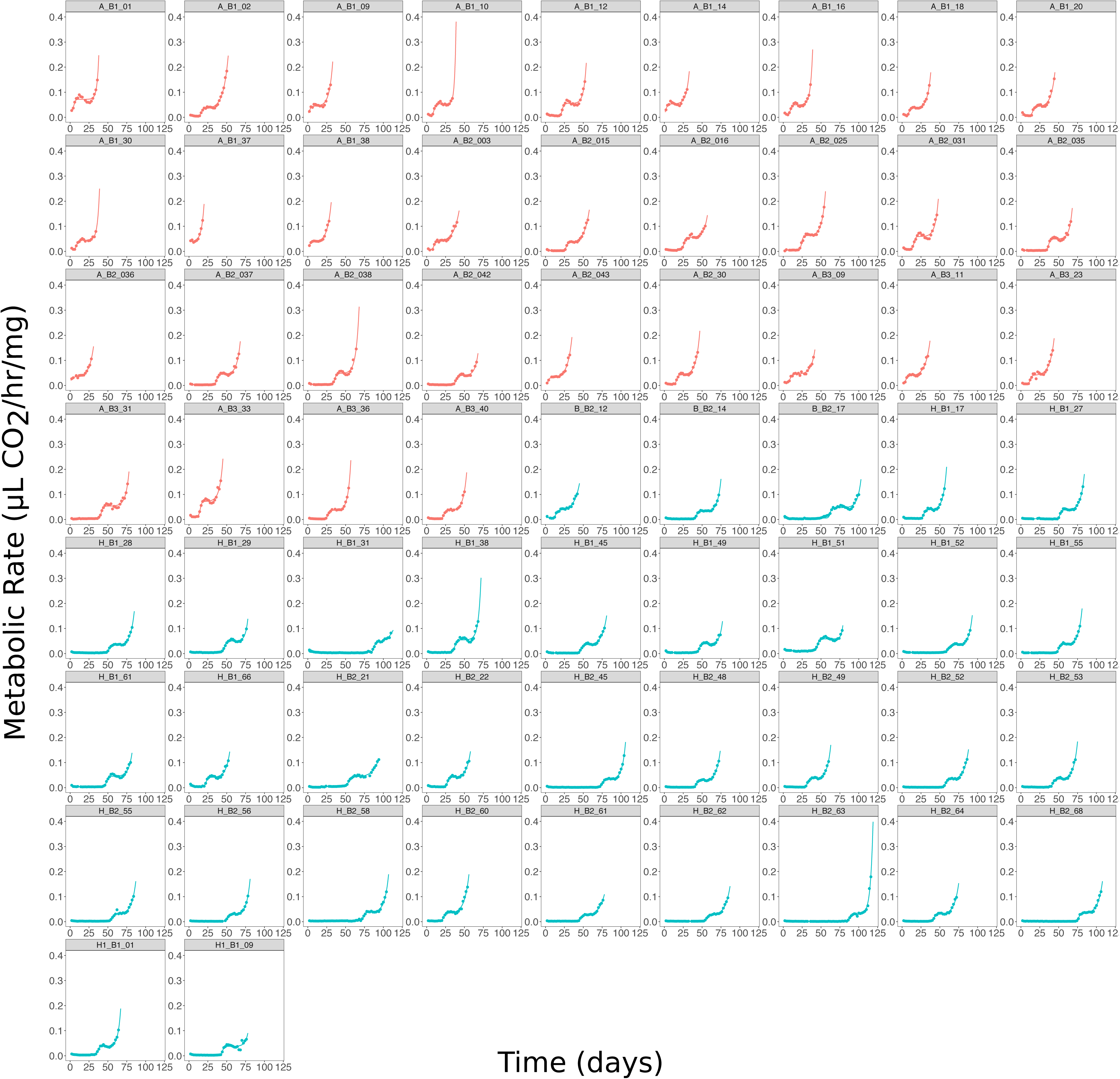
Plots of mass adjusted metabolic rates at each time point along with curves fitted for the best-fit non-linear metabolic trajectories for each individual apple fly (orange points) and hawthorn fly (blue points) in experiment 1.

**Figure S2.**
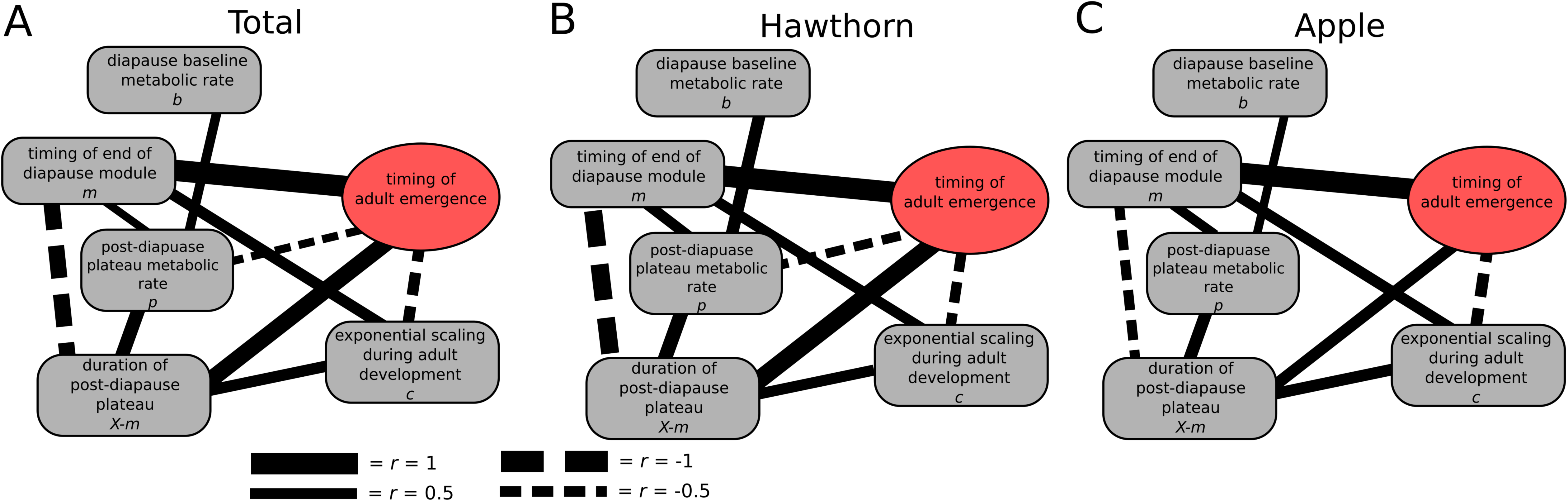
Network of significant (P < 0.05) partial correlations between each fitted parameter, representing a different component of diapause development, from the best-fit non-linear models of metabolic trajectories and adult eclosion time for A) all flies, B) hawthorn flies, and C) apple flies from experiment 1, showing that partial correlation relationships among parameters and emergence timing remain largely the same between both host races. Line weights are proportional to the magnitude of Pearson’s *r*, for both positive correlations (solid lines) and negative correlations (dashed lines).

**Figure S3.**
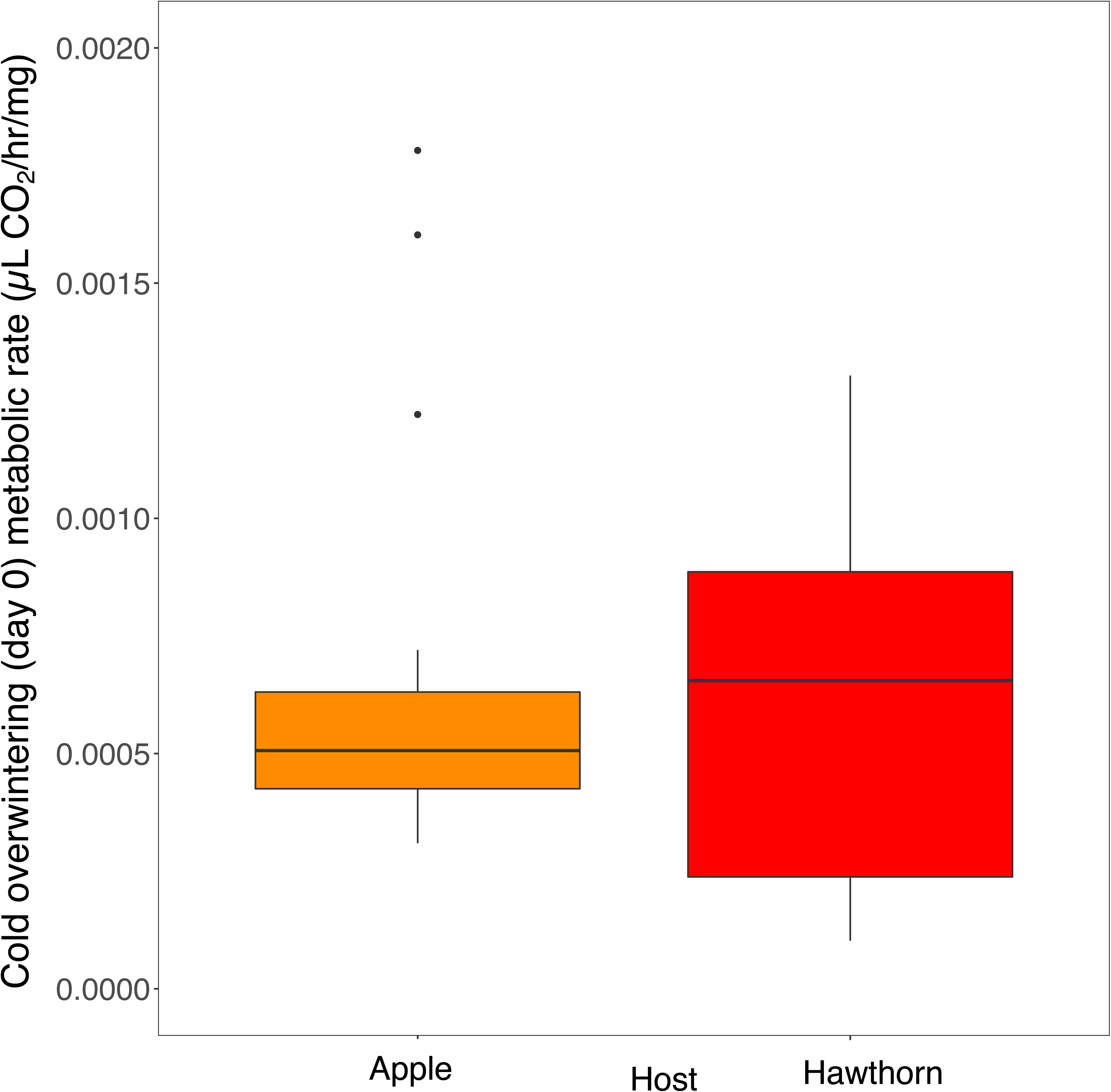
Box plot showing that mean overwintering metabolic rates did not differ between apple flies (yellow) and hawthorn flies (red) from experiment 2. Values come from the measurement of flies over a 24-hour interval during continuous simulating overwintering conditions prior to their removal from the cold. These data are represented as time point zero in Figure 5.

**Figure S4.**
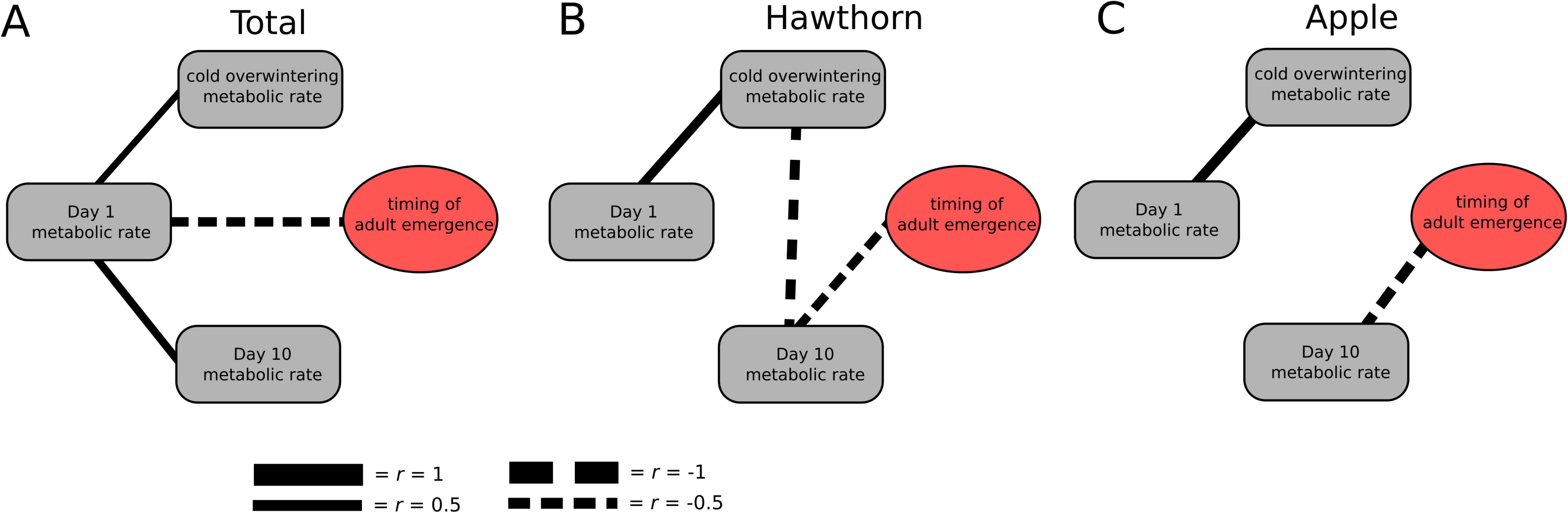
Scatter plots showing A) the significant inverse relationship between *(r* = −0.318) between day 1 metabolic rate and adult emergence time, with higher initial metabolic rate peaks leading to earlier emergence times and B) the non-significant relationship between day 10 metabolic rate and adult emergence timing, with three outlier individuals that have likely exited the diapause maintenance phase by day 10 for each apple fly (yellow) and hawthorn fly (red) in experiment 2.

**Figure S5.**
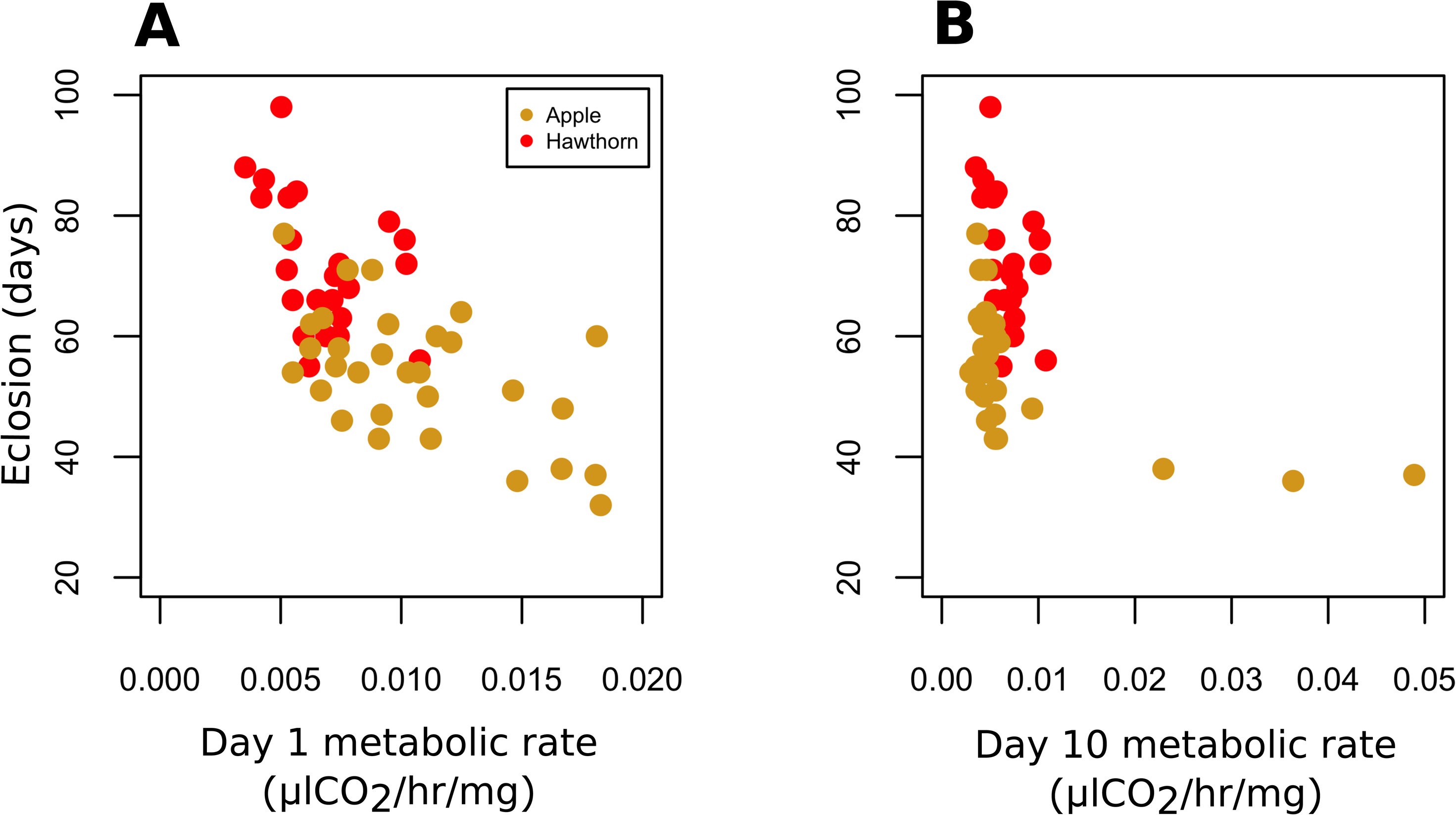
Networks of significant (P < 0.05) partial correlations between overwintering metabolic rate, day 1 metabolic rate (reflective of the initial post-winter metabolic increase), and day 10 metabolic rates (reflective of post-winter diapause baseline metabolic rates) and adult eclosion for A) all flies, B) hawthorn flies, and C) apple flies from experiment 2. Line weights are proportional to the magnitude of Pearson’s *r*, for both positive correlations (solid lines) and negative correlations (dashed lines).

**Table S1.** AICc values for each of the three hierarchical non-linear models of metabolic trajectories for each fly in experiment 1.

| Fly | Host Race | AICc values |  |  |
| --- | --- | --- | --- | --- |
|  |  | monophasic | biphasic | triphasic |
| A1 | Apple | -85.42488 | -101.16601 | -95.87575 |
| A2 | Apple | -159.3352 | -195.2199 | -192.0269 |
| A3 | Apple | -90.23488 | -93.30185 | -86.33917 |
| A4 | Apple | -137.7368 | -183.5841 | -182.2060 |
| A5 | Apple | -87.73685 | -98.39063 | -91.77090 |
| A6 | Apple | -100.4272 | -133.4496 | -128.6462 |
| A7 | Apple | -116.0277 | -154.3354 | -151.7218 |
| A8 | Apple | -125.1148 | -149.7043 | -145.7560 |
| A9 | Apple | -96.47984 | -126.50280 | -120.66406 |
| A10 | Apple | -68.01215 | -54.19323 | -13.90317 |
| A11 | Apple | -96.65276 | -99.46503 | -92.12180 |
| A12 | Apple | -120.4935 | -134.1151 | -129.5280 |
| A13 | Apple | -185.0641 | -222.9163 | -221.6227 |
| A14 | Apple | -132.8936 | -164.5961 | -161.3035 |
| A15 | Apple | -117.2497 | -146.7242 | -142.3276 |
| A16 | Apple | -188.1717 | -246.0007 | -247.0824 |
| A17 | Apple | -96.76512 | -101.30425 | -93.07224 |
| A18 | Apple | -175.6545 | -221.7109 | -218.5474 |
| A19 | Apple | -99.23066 | -110.27054 | -103.61684 |
| A20 | Apple | -107.6863 | -118.5794 | -113.3541 |
| A21 | Apple | -126.1644 | -145.1380 | -141.5845 |

Table S1 continued
| Fly | Host Race | AICc values |  |  |
| --- | --- | --- | --- | --- |
|  |  | monophasic | biphasic | triphasic |
| A22 | Apple | -217.5604 | -291.0552 | -288.7704 |
| A23 | Apple | -167.1777 | -240.5444 | -240.7968 |
| A24 | Apple | -157.1383 | -201.4665 | -199.8782 |
| A25 | Apple | -104.9526 | -136.0942 | -132.2242 |
| A26 | Apple | -140.7678 | -207.0548 | -203.6812 |
| A27 | Apple | -198.7765 | -258.1858 | -256.7039 |
| A28 | Apple | -156.5090 | -213.0256 | -210.2791 |
| A29 | Apple | -92.64481 | -127.87155 | -122.72742 |
| A30 | Apple | -117.6916 | -144.5089 | -140.4335 |
| A31 | Apple | -214.7300 | -275.9143 | -275.9728 |
| H1 | Hawthorn | -214.3493 | -285.7186 | -288.1658 |
| H2 | Hawthorn | -238.7159 | -282.3931 | -281.0323 |
| H3 | Hawthorn | -180.8670 | -233.9812 | -231.9984 |
| H4 | Hawthorn | -259.3621 | -351.8553 | -354.2683 |
| H5 | Hawthorn | -292.7671 | -386.2696 | -386.8440 |
| H6 | Hawthorn | -395.1055 | -476.3699 | -499.9463 |
| H7 | Hawthorn | -192.0993 | -250.6646 | -248.1271 |
| H8 | Hawthorn | -265.4327 | -326.4221 | -327.5829 |
| H9 | Hawthorn | -240.5597 | -293.8475 | -297.6247 |
| H10 | Hawthorn | -326.6583 | -408.7803 | -410.9211 |

Table S1 continued
| Fly | Host Race | AICc values |  |  |
| --- | --- | --- | --- | --- |
|  |  | monophasic | biphasic | triphasic |
| H11 | Hawthorn | -265.2983 | -353.9769 | -358.0998 |
| H12 | Hawthorn | -244.5326 | -308.4879 | -310.6853 |
| H13 | Hawthorn | -161.4259 | -195.7099 | -194.0092 |
| H14 | Hawthorn | -133.3469 | -155.4459 | -153.0507 |
| H15 | Hawthorn | -254.4221 | -343.1827 | -354.8459 |
| H16 | Hawthorn | -318.0212 | -382.7139 | -381.1767 |
| H17 | Hawthorn | -177.9566 | -225.7985 | -222.3016 |
| H18 | Hawthorn | -376.5118 | -504.9609 | -513.9705 |
| H19 | Hawthorn | -261.0556 | -337.7621 | -338.1983 |
| H20 | Hawthorn | -203.6956 | -269.0703 | -265.8444 |
| H21 | Hawthorn | -315.5219 | -386.2596 | -387.6650 |
| H22 | Hawthorn | -246.0133 | -303.8118 | -302.6023 |
| H23 | Hawthorn | -304.7557 | -359.4706 | -357.3332 |
| H24 | Hawthorn | -287.6338 | -397.2828 | -399.9917 |
| H25 | Hawthorn | -376.0554 | -474.7933 | -472.3875 |
| H26 | Hawthorn | -171.2786 | -204.8060 | -201.5091 |
| H27 | Hawthorn | -301.8461 | -350.0046 | -350.3595 |
| H28 | Hawthorn | -327.8362 | -396.3005 | -399.3649 |
| H29 | Hawthorn | -383.5556 | -482.3031 | -480.4219 |
| H30 | Hawthorn | -259.9859 | -315.4500 | -314.0465 |
| H31 | Hawthorn | -395.3495 | -493.7767 | -494.3287 |

Table S1 continued
|  |  | **AICc values** |  |  |
| --- | --- | --- | --- | --- |
| **Fly** | **Host Race** | **monophasic** | **biphasic** | **triphasic** |
| H32 | Hawthorn | -233.9627 | -311.3159 | -315.6969 |
| H33 | Hawthorn | -223.9982 | -308.7444 | -311.5882 |
| H34 | Hawthorn | -292.5576 | -372.8700 | -370.3306 |
| H35 | Hawthorn | -217.4432 | -267.6473 | -265.6989 |

**Table S2.** Results of paired t-tests between parameter coefficients from biphasic and triphasic trajectory models from the same fly.

| Parameter | Description | *t* | p-value | mean of differences |
| --- | --- | --- | --- | --- |
| *b* | Baseline metabolic rate | 1.760 | 0.083 | 0.00104 |
| *p* | Plateau metabolic rate | -0.827 | 0.411 | -0.00021 |
| *m* | Timing of diapause exit | 1.152 | 0.254 | 0.11743 |
| *X* | Timing of exponential increase | -0.756 | 0.453 | -0.07246 |
| *c* | Exponential scaling | -0.639 | 0.525 | -0.00088 |

**Table S3.** Summary of repeated-measured ANOVA analysis of metabolic rate over time as a function of host race.

|  | **Df** | **SS** | **MS** | **F-value** | **p-value** |
| --- | --- | --- | --- | --- | --- |
| *Host* | 1 | 1985 | 1985 | 21.762 | <0.001 |
| *Day 1* | 1 | 2260 | 2260.1 | 24.779 | <0.001 |
| *Day 9* | 1 | 53 | 52.8 | 0.578 | 0.4515 |
| *Day 10* | 1 | 120 | 120.2 | 1.318 | 0.2579 |
| *Day 11* | 1 | 17 | 17.1 | 0.188 | 0.667 |
| *Host x day 1* | 1 | 429 | 428.9 | 4.702 | <0.05 |
| *Host x day 9* | 1 | 216 | 216 | 2.368 | 0.1319 |
| *Host x day 10* | 1 | 564 | 563.9 | 2.182 | <0.05 |
| *Residuals* | 39 | 3557 | 91.2 |  |  |

